# Topology-Based Query Framework for Longitudinal Omics Trajectories

**DOI:** 10.64898/2026.08.18.745427

**Authors:** Nazanin Zounemat-Kermani, Matthew Richardson, Alen Faiz, Siyao Wang, Kai Sun, Dragana Vuckovic, Maarten van den Berge, Anke-Hilse Maitland-van der Zee, Ian Sayers, Sven-Erik Dahlén, Christopher Brightling, Salman Siddiqui, Kian Fan Chung, Martijn C. Nawijn, Marc Chadeau-Hyam, Ian Michael Adcock

**Author notes:** Corresponding author. August 18, 2026.

## Abstract

1

**Background:** Many longitudinal omics studies contain only a small number of repeated measurements collected before, during, or after an intervention. Existing approaches, including mixed-effects models and generalized additive models, estimate temporal effects but do not generally provide a discrete representation of trajectory topology that can be queried directly across experimental groups.

**Methods:** We developed LongOmicsTraj, an open-source R package for topology-based representation and querying of short longitudinal omics trajectories. The framework encodes the direction of change between adjacent visits as *up*, *down*, or *flat*, with the ordered sequence defining an Ordinal Trajectory State (OTS). LongOmicsTraj operates downstream of trajectory estimation and can therefore be applied to empirical summaries or model-derived visit-level estimates, including those from linear mixed-effects models, generalized additive models, and polynomial regression, following a maSigPro-style time-course formulation [1]. OTS labels provide a common representation for topology-based querying, cross-group comparison, and evaluation of higher-level representations such as trajectory clusters. We evaluated the framework using controlled simulations and bronchial biopsy transcriptomic data from the GLUCOLD corticosteroid intervention study (GEO accession `GSE36221`), measured at baseline, 6 months, and 30 months. The biological analysis compared continued inhaled corticosteroid (ICS) treatment, ICS withdrawal after 6 months, and placebo.

**Results:** In simulations, LongOmicsTraj recovered predefined stable, monotonic, transient, rebound, and oscillatory trajectories with high accuracy when longitudinal signal was sufficiently clear, with performance declining under high-noise conditions and depending partly on the upstream estimator. In GLUCOLD, comparator-aware topology queries reduced 20,358 measured transcripts to 168 genes showing a corticosteroid response that was maintained during continued treatment, reversed following withdrawal, and was not reproduced under placebo. The selected genes included established corticosteroid-response genes and were enriched for immune-cell migration, chemotaxis, cell adhesion, and extracellular-matrix organisation. Topology-aware evaluation of FlexMix trajectory clusters additionally revealed substantial within-cluster temporal heterogeneity, with topology purities of approximately 46–60%.

**Conclusions:** LongOmicsTraj provides a compact, directly queryable representation of temporal direction and order in short longitudinal omics studies. It complements existing longitudinal estimation and clustering methods by making trajectory structure explicit, enabling structured cross-group queries and quantification of temporal heterogeneity within trajectory clusters.

## 2 Introduction

Longitudinal omics studies are increasingly used in respiratory disease research to support precision medicine. Repeated sampling of the same individuals provides an opportunity to characterise disease trajectories and their stability over time, distinguishing persistent biological states from treatment response, recovery, progression and relapse. Transcriptomic, proteomic, metabolomic and imaging measurements are therefore increasingly collected across sequential visits. Large longitudinal studies, including GLUCOLD, PRISM, APEX and 3TR, have been established to investigate how airway biology evolves over time and in response to therapy [2, 3, 4, 5, 6].

A central analytical challenge is how to represent temporal behaviour while preserving its biological meaning. Longitudinal analyses commonly use empirical visit summaries or model-based estimates from linear mixed-effects models, generalized additive models and polynomial regression [7, 8, 1]. These approaches answer complementary statistical questions: empirical summaries describe observed visit-level changes, mixed-effects models account for repeated measurements, generalized additive models capture smooth non-linear responses, and polynomial regression represents structured temporal trends. However, their outputs do not usually encode the direction and order of temporal change as a discrete, directly queryable object. Biologically distinct behaviours—including sustained response, delayed activation, transient suppression, rebound and oscillation—may therefore have similar endpoint differences despite reflecting different temporal processes (Figure 1).

**Figure 1:**
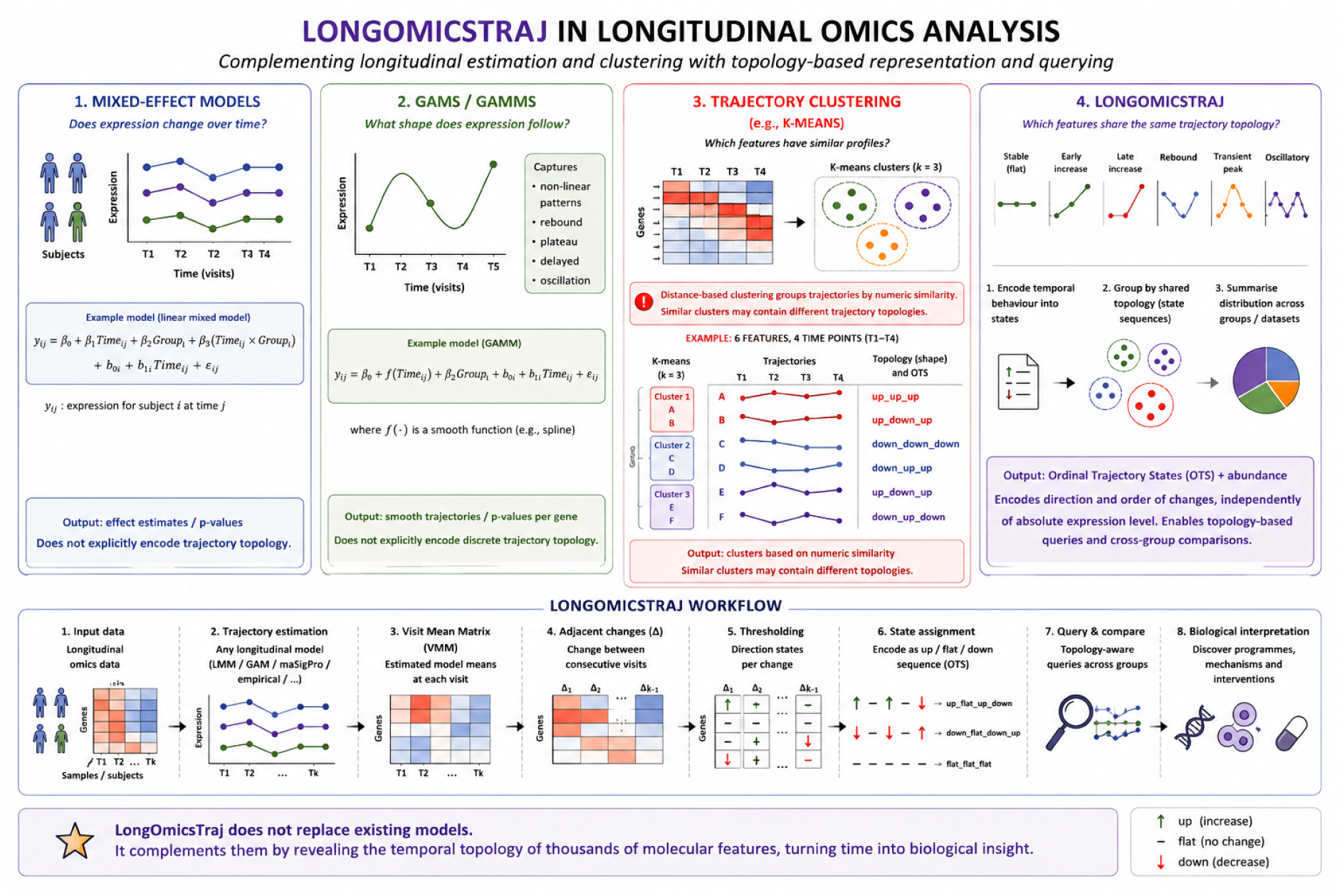
Overview of LongOmicsTraj. Empirical summaries and longitudinal models estimate temporal behaviour, whereas LongOmicsTraj represents the direction and order of change as discrete Ordinal Trajectory States (OTS). Visit-level estimates are converted into adjacent changes, thresholded states and OTS labels for topology-based querying, cross-group comparison and biological interpretation. LongOmicsTraj complements, rather than replaces, existing longitudinal estimators.

This distinction is particularly relevant to studies with relatively few clinically meaningful visits. Respiratory cohorts often contain only three to five time points because procedures such as bronchoscopy, sputum induction and repeated imaging are invasive, costly or difficult to repeat. In such settings, each intermediate transition can carry substantial information. For example, sustained suppression and suppression followed by rebound may have similar baseline-to-final differences but different intermediate trajectories. Missing observations add a further complication: mixed-effects and related model-based estimators can accommodate unbalanced repeated measurements under their modelling assumptions, whereas empirical summaries require sufficient observations at each visit.

LongOmicsTraj was developed to address this representational problem. Rather than replacing longitudinal estimators, it provides a topology layer that operates on their visit-level estimates (Figure 1). For each molecular feature, changes between adjacent visits are classified as *up*, *down* or *flat* and concatenated into an Ordinal Trajectory State (OTS). A gene that decreases between baseline and 6 months and increases between 6 and 30 months, for example, is represented as `down_up`. Because trajectory estimation and topology assignment are separated, the same representation can be applied to empirical summaries or estimates obtained from mixed-effects models, generalized additive models or polynomial regression.

The resulting representation retains the order and direction of temporal change while remaining compact and queryable. It supports structured comparisons across experimental groups and can distinguish different phases of treatment response, for example, an early acute response from a later sustained response. It also enables identification of features that rebound after treatment withdrawal but remain suppressed under continued therapy, grouping of features with shared topologies for downstream enrichment analysis, and comparison of trajectory-state distributions between biological or treatment groups. The same framework can also be applied to higher-level representations, including trajectory clusters and module-level profiles.

We illustrate LongOmicsTraj using the GLUCOLD corticosteroid intervention study in chronic obstructive pulmonary disease, with bronchial biopsy transcriptomic measurements at baseline, 6 months and 30 months [2, 3]. The biological analysis focuses on continued in-haled corticosteroid (ICS) treatment, ICS withdrawal after 6 months, and placebo; the smaller ICS/LABA (inhaled corticosteroid/long-acting *β*_2_-agonist) group was omitted to simplify biological interpretation. GLUCOLD provides a useful case study because corticosteroid treatment and withdrawal can generate non-monotonic molecular responses, including suppression, rebound and delayed reversal, that are not fully described by endpoint comparisons alone.

Here, we present LongOmicsTraj and evaluate it using controlled simulations and the GLUCOLD longitudinal intervention study. We assess whether topology-based representation can recover predefined temporal patterns, distinguish trajectories with similar endpoint behaviour but different intermediate dynamics, support structured cross-group queries, and provide an interpretable complement to existing longitudinal estimators.

## 3 Materials and methods

### 3.1 LongOmicsTraj framework

LongOmicsTraj separates trajectory estimation from topology representation. The framework proceeds from visit-level trajectory estimates to adjacent temporal changes, threshold-based ordinal state assignment, Ordinal Trajectory State (OTS) construction, and topology-based querying (Figure 1). Each stage is described below.

### 3.2 Visit Mean Matrix

LongOmicsTraj separates the estimation of longitudinal trajectories from their representation. The input to the framework is a *Visit Mean Matrix* (VMM), which stores visit-level estimates for each longitudinal object. An object may represent a molecular feature (e.g. gene, protein or metabolite), a pathway score, a module eigengene, a cell population, or any higher-level longitudinal representation.

Let *i* = 1*, . . ., N* index longitudinal objects and *t* = 1*, . . ., T* denote ordered study visits.

For each object, an upstream longitudinal analysis produces the visit-level estimate

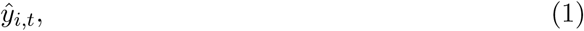

where *ŷ_i,t_* denotes the estimated value of object *i* at visit *t*.

The VMM is intentionally estimator-agnostic and may be constructed from empirical visit means or from any longitudinal modelling framework that produces visit-level estimates, including linear mixed-effects models, generalised additive models and other regression-based approaches [7, 8]. Model-derived VMMs represent covariate-adjusted estimates of the expected value at each visit, whereas empirical VMMs summarise the observed average trajectory. The downstream topology framework is not tied to a particular statistical estimator; however, the resulting topology assignments depend on the visit-level estimates supplied to the framework.

When multiple experimental groups are analysed, the notation extends naturally to

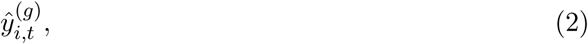

where *g* indexes the experimental group. All subsequent quantities are computed independently within each group.

### 3.3 Adjacent temporal changes

For each object and pair of adjacent visits,

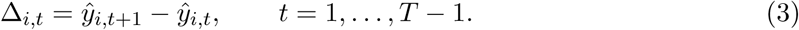

The ordered sequence of adjacent changes summarizes the temporal evolution of each object and provides a common representation across all upstream estimation methods.

For studies containing multiple experimental groups,

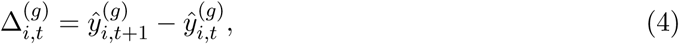

is computed independently within each group.

The VMM is assumed to contain complete visit-level estimates. When trajectory estimators such as linear mixed-effects models or generalized additive models are used, incomplete longitudinal observations are accommodated during trajectory estimation, whereas empirical summaries require complete or preprocessed visit-level data.

### 3.4 Adaptive thresholding

Each adjacent change is classified relative to a threshold representing the minimum biologically meaningful change. The objective is to distinguish meaningful temporal changes from background variability rather than to assess statistical significance.

LongOmicsTraj provides five thresholding strategies.

#### Fixed threshold

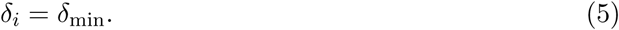

#### Global MAD

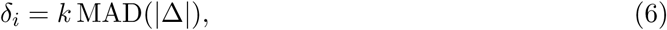

where the threshold is estimated from the median absolute deviation of all adjacent changes.

#### Object-specific MAD

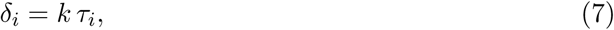

where *τ_i_*denotes the median absolute deviation of the adjacent changes for object *i*.

#### Hybrid threshold (default)

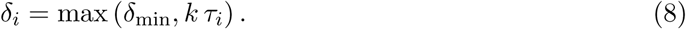

This combines a minimum effect-size threshold with an object-specific variability estimate.

#### Quantile threshold

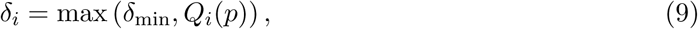

where *Q_i_*(*p*) denotes the *p*th quantile of the absolute adjacent changes for object *i*.

The parameters *δ*_min_, *k*, and *p* are user configurable. When not specified, *δ*_min_ is initialized from the empirical distribution of adjacent changes and can subsequently be refined through sensitivity analysis.

### 3.5 Ordinal state assignment

Each adjacent change is mapped to one of three ordinal states,

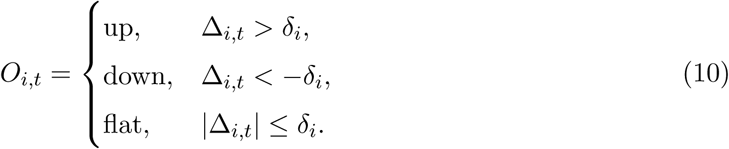

When multiple groups are analysed, the notation extends to *O_i,t_*^(*g*)^.

#### 3.5.1 Choosing threshold parameters

The thresholding parameters determine the minimum adjacent change classified as biologically meaningful. Their appropriate values depend on the measurement platform, biological variability and scientific objective. Consequently, LongOmicsTraj does not recommend a universally optimal thresholding strategy.

All thresholding methods are fully user configurable. The hybrid threshold (Equation 8) is used as the package default because it combines a minimum effect-size criterion with an object-specific estimate of variability. This and all other parameter defaults are implementation choices provided for convenience rather than methodological recommendations. Users are encouraged to assess the sensitivity of their results to alternative thresholding strategies and parameter values.

### 3.6 Ordinal Trajectory States

The ordered sequence of ordinal states defines the *Ordinal Trajectory State* (OTS),

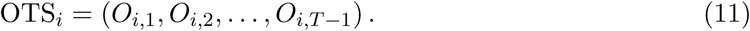

For convenience, the sequence is represented as a topology string obtained by concatenating adjacent states. For example,

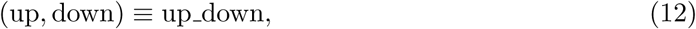

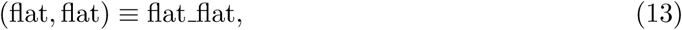

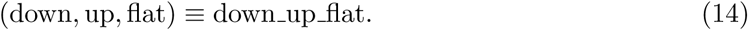

For studies containing multiple experimental groups, the notation extends naturally to

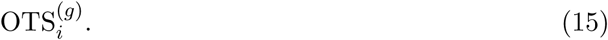

With *T* visits, each object is represented by *T −* 1 ordinal states and a single OTS.

### 3.7 Statistical support

LongOmicsTraj optionally stores statistical support for each adjacent transition. For transition

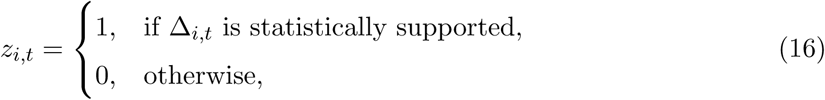

where statistical support may be obtained from any upstream hypothesis-testing procedure. These indicators are retained alongside the corresponding OTS but do not influence topology assignment. Consequently, trajectory representation remains independent of statistical significance while allowing statistical evidence to be incorporated during downstream analyses.

### 3.8 Topology-based queries

LongOmicsTraj identifies temporal patterns by querying one or more OTS. For studies containing multiple experimental groups, queries specify allowable topology strings for each group and return all objects satisfying those constraints.

For example,

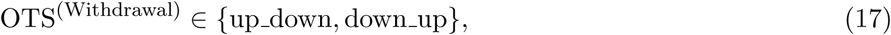

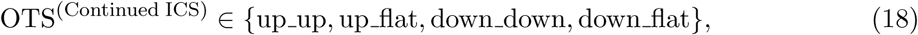

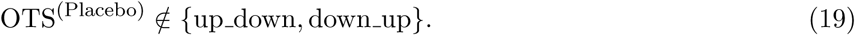

This query identifies objects exhibiting reversal following treatment withdrawal while excluding comparable placebo trajectories.

Because querying operates on the resulting OTS labels, the same query definitions apply regardless of the estimator used to construct the VMM. Estimation, representation and querying therefore remain distinct components of the analytical framework. Estimation, representation, and querying therefore remain distinct components of the analytical framework.

### 3.9 Extension to higher-level representations

LongOmicsTraj is applicable to any longitudinal object represented by a sequence of visit-level estimates. In addition to individual molecular features, this includes pathway scores, module eigengenes, latent variables, cell populations, cluster centroids and other derived longitudinal representations.

For any representation *r*, the adjacent changes, thresholding procedure, ordinal state assignment and OTS construction are identical to those defined above,

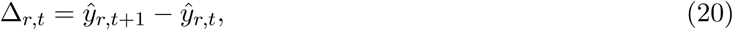

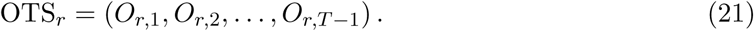

Consequently, topology-based queries operate within a common discrete trajectory space irrespective of the level of biological organization, allowing direct comparison of temporal patterns across genes, pathways, clusters and other higher-level representations.

## 4 Software implementation

LongOmicsTraj is implemented as an open-source R package using an S4 object-oriented architecture. The central `LongOmicsTraj` object stores the complete longitudinal analysis, including the input `MultiAssayExperiment`, the Visit Mean Matrix (VMM), adjacent changes, adaptive thresholds, Ordinal Trajectory States (OTS), relationships between biological representations (e.g. genes, pathways and clusters), analysis results and analysis metadata. This design provides a unified representation of longitudinal analyses while maintaining full provenance throughout the workflow.

The analysis pipeline is organised into independent stages. Visit-level estimates are first computed using an upstream estimator and stored in the VMM. Adjacent temporal changes are then calculated, feature-specific thresholds are estimated, and OTS are assigned. This modular design allows LongOmicsTraj to operate on outputs from multiple longitudinal estimation procedures, including empirical visit means, linear mixed-effects models [7], generalized additive models [8], and maSigPro [1], while producing a common topology representation for downstream querying and interpretation. Additional estimators can be incorporated without modifying the downstream topology framework.

LongOmicsTraj accepts multi-omics data through the `MultiAssayExperiment` framework [9], allowing multiple molecular assays to be stored within a single object while preserving sample alignment and associated metadata. The resulting S4 object provides a common interface for topology assignment, querying, clustering, enrichment analysis and visualisation across multiple omics layers.

The package is currently under review for inclusion in Bioconductor. The development version is freely available from GitHub.

https://github.com/nzkermani/LongOmicsTraj

and can be installed using

~~~
devtools::install_github("nzkermani/LongOmicsTraj")
~~~

A typical analysis consists of five sequential steps:

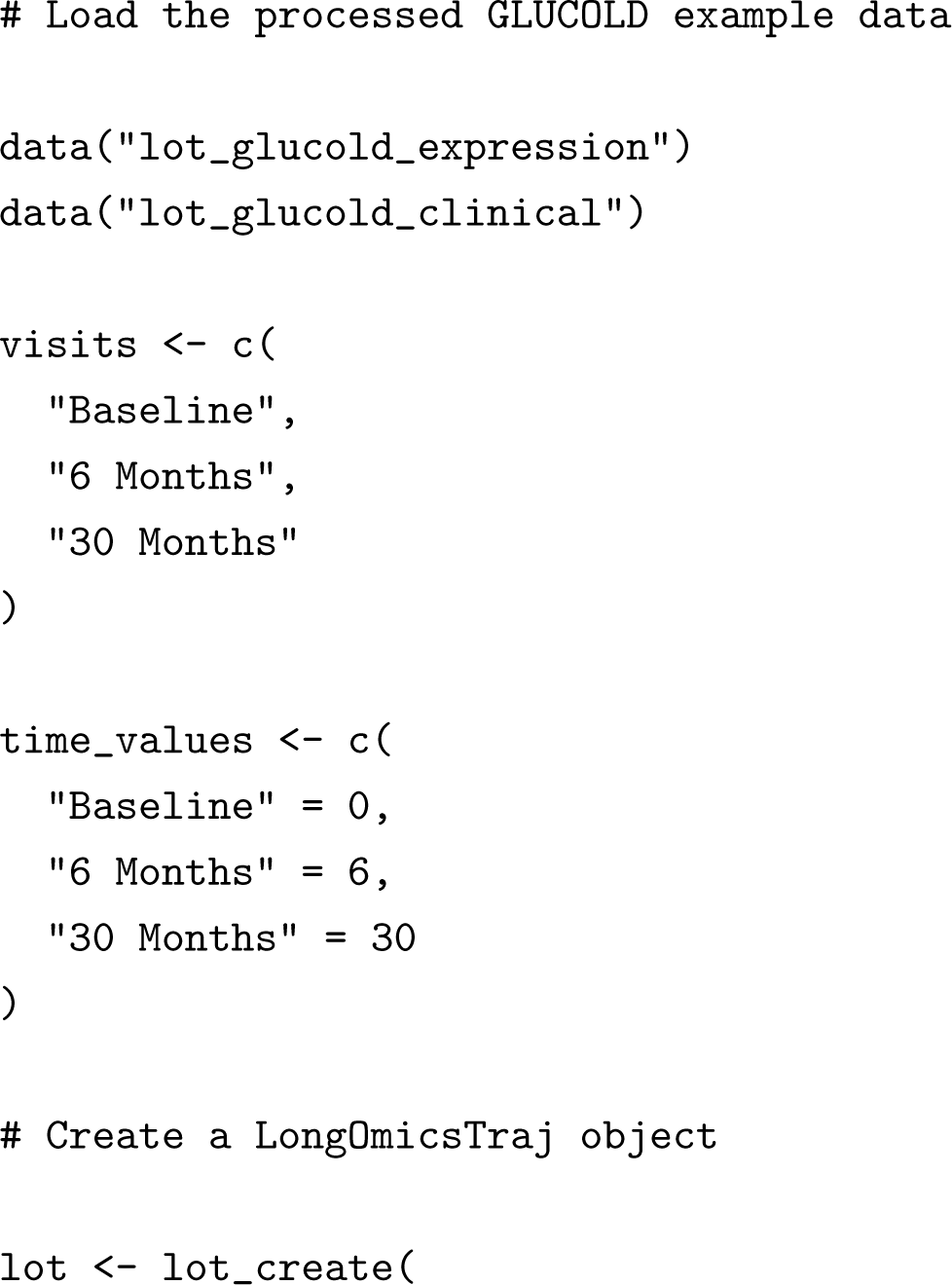
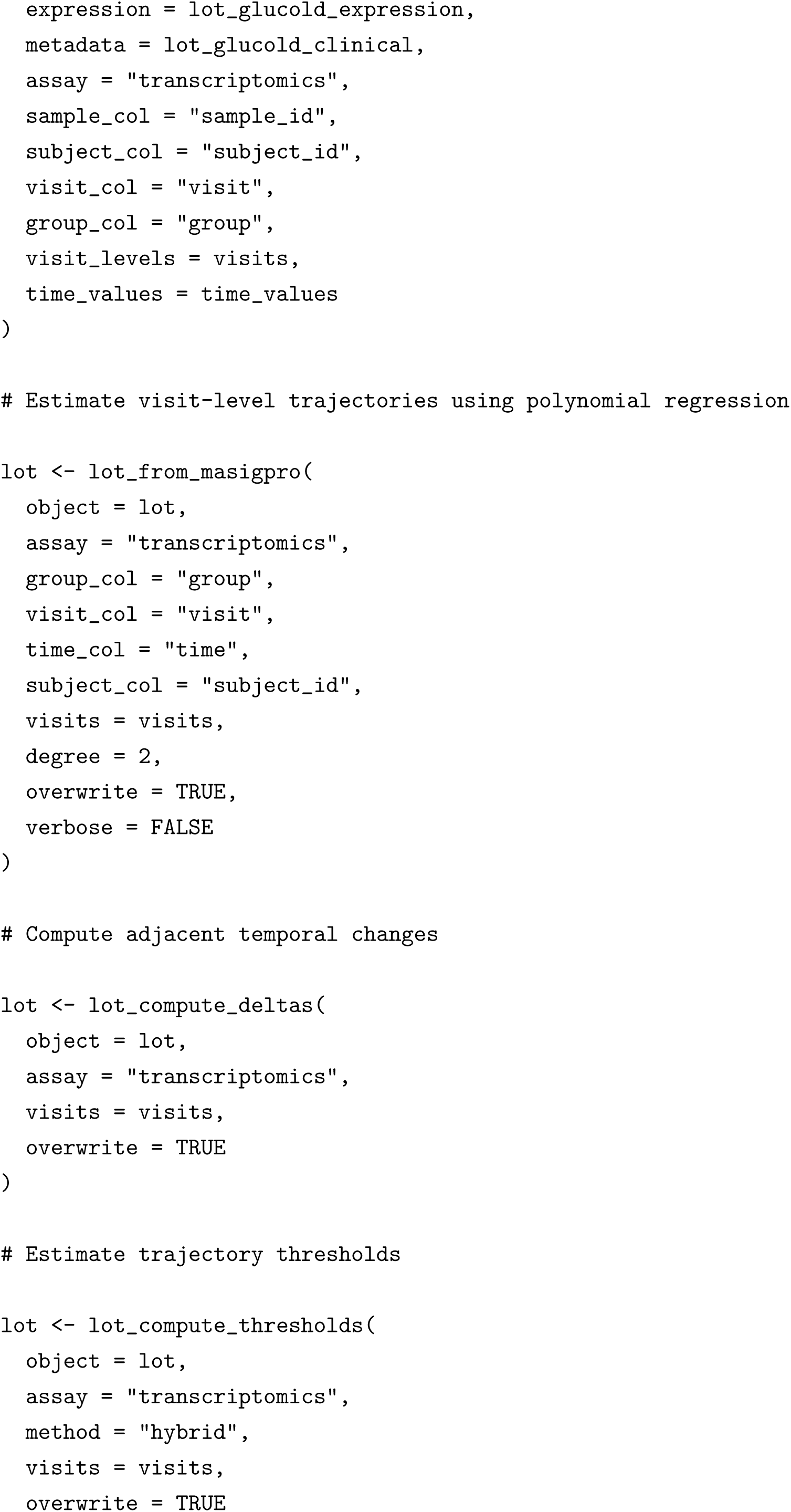
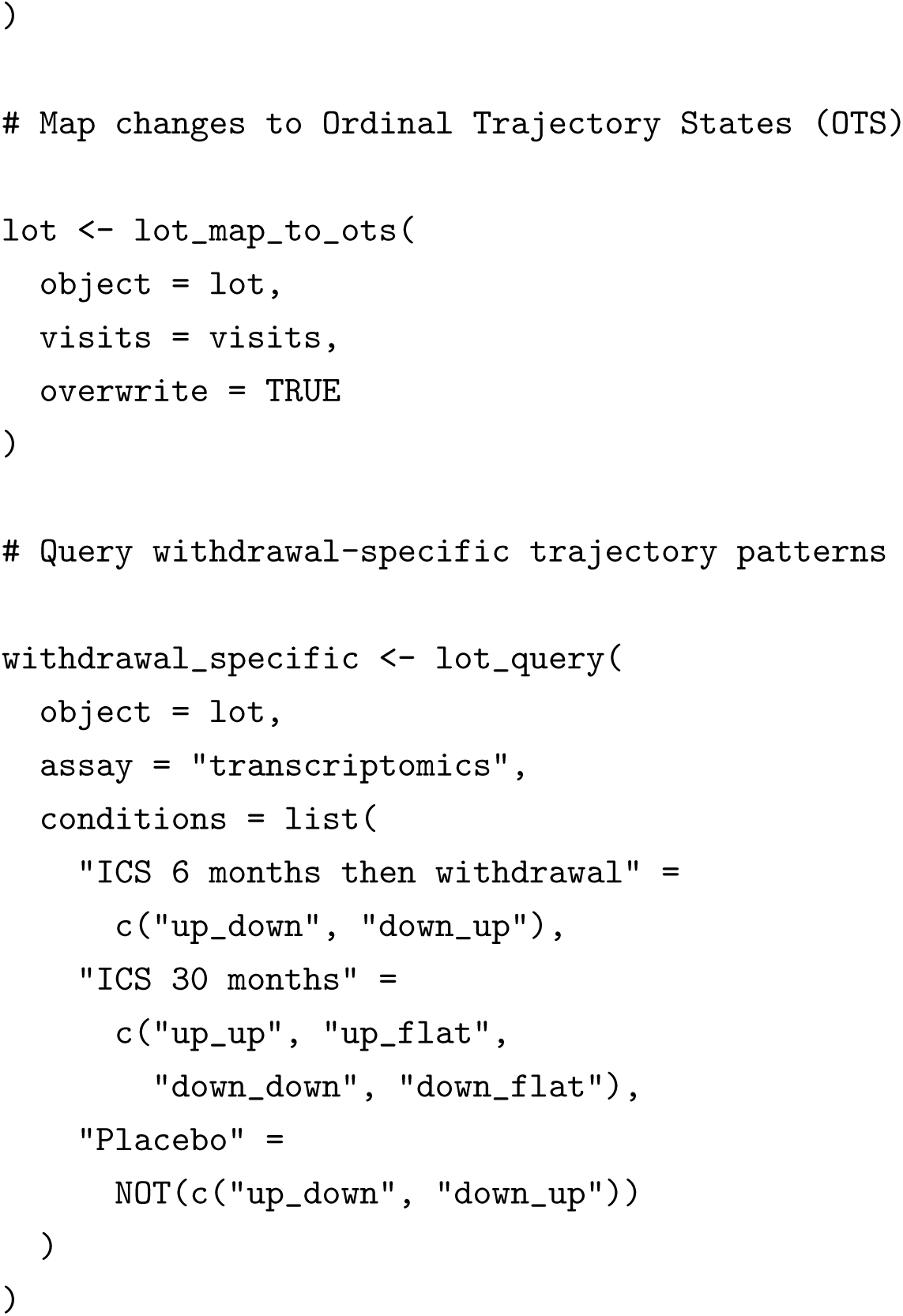

Additional methods support topology-based querying, trajectory clustering, higher-level representations, estimator comparison and visualisation.

## 5 Simulation study and real-world application

### 5.1 Simulation study

We used simulation to test whether LongOmicsTraj could recover longitudinal patterns when the true trajectory was known. Synthetic datasets contained 900 molecular features measured at four visits in 30 subjects. For feature *i*, subject *j*, and visit *t*, observations were generated as

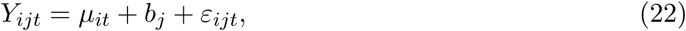

where *µ_it_* is the predefined mean trajectory, *b_j_ ∼ N* (0, *σ*_b_^2^) represents between-subject variability, and *ε_ijt_∼ N* (0*, σ*^2^) represents random variation around the underlying trajectory.

The predefined trajectories were constructed from combinations of upward, downward and stable changes between consecutive visits. The effect-size parameter controlled the magnitude of these true visit-to-visit changes: small values produced subtle temporal responses, whereas large values produced clearly separated trajectories. The noise parameter controlled how much the observed measurements varied around the true trajectory, representing biological and measurement variability. Thus, a small effect combined with high noise represented a difficult setting in which a genuine temporal response was weak relative to background variation, whereas a large effect with low noise represented a clearly detectable trajectory.

We considered three effect sizes, *{*0.4,;1.2,;2.4*}*, and three noise levels *{*0.2,;0.4,;0.8*}*. These values were chosen to span low, intermediate and high signal and variability conditions rather than to represent specific clinical effect sizes. Their factorial combination produced nine signal–noise settings. Each setting was independently replicated 30 times, giving 3 × 3 × 30 = 270 simulated datasets. Within each dataset, features followed predefined stable, monotonic, rebound, oscillatory and mixed trajectory patterns. Because the true topology of every feature was known by construction, we could directly measure how often LongOmicsTraj recovered the correct temporal pattern.

The simulations were designed to answer three practical questions: whether LongOmicsTraj recovers the correct topology when longitudinal signal is clear; how recovery deteriorates as temporal changes become smaller or measurements become noisier; and whether performance is consistent when visit-level trajectories are estimated using different longitudinal methods. We additionally recorded computational time. This provides a controlled assessment of the conditions under which topology-based representation is reliable and where caution is required before applying it to biological data.

### 5.2 GLUCOLD study and transcriptomic data

LongOmicsTraj was applied to publicly available bronchial biopsy transcriptomic data from the GLUCOLD randomized clinical trial [2, 3]. Gene-expression data and sample metadata were obtained from the NCBI Gene Expression Omnibus (GEO; accession GSE36221) and were generated using the Affymetrix Human Gene 1.0 ST Array (GPL6244). Samples were collected at baseline, 6 months and 30 months. The available dataset included participants assigned to placebo, continued inhaled corticosteroid (ICS), ICS withdrawal after 6 months, or continued ICS/LABA treatment. Participant characteristics shown in Table 1 were derived from the publicly available GEO sample metadata. For the topology-based biological analysis, we focused on the placebo, continued ICS and ICS-withdrawal groups; the smaller ICS/LABA group was omitted to simplify interpretation of treatment maintenance and withdrawal. Detailed descriptions of the GLUCOLD study design, recruitment criteria, treatment allocation and clinical characteristics have been reported previously [2, 3].

**Table 1:** Participant characteristics in the GLUCOLD transcriptomic dataset. Age and sex were summarised once per unique participant. Values are median (minimum–maximum) for age and number (%) for sex.

| Treatment group | N | Age, median (min–max) | Female, n (%) | Male, n (%) |
| --- | --- | --- | --- | --- |
| Placebo | 23 | 61.0 (46.0–73.0) | 4 (17.4%) | 19 (82.6%) |
| ICS 30 months | 25 | 61.0 (47.0–74.0) | 3 (12.0%) | 22 (88.0%) |
| ICS 6 months then withdrawal | 22 | 65.5 (47.0–74.0) | 3 (13.6%) | 19 (86.4%) |
| ICS/LABA 30 months | 20 | 58.5 (45.0–74.0) | 3 (15.0%) | 17 (85.0%) |

Data import was performed in R using the Bioconductor package `GEOquery`[10]. For each treatment group, visit-level estimates were generated using the longitudinal estimators described above and stored in a common Visit Mean Matrix (VMM). Adjacent changes were then mapped to Ordinal Trajectory States (OTS), and topology-based queries were used to identify temporal patterns specific to corticosteroid withdrawal.

### 5.3 Topology-aware trajectory clustering

LongOmicsTraj extends trajectory estimation by providing topology-aware clustering of longitudinal molecular profiles. The objective is to group molecular features with similar temporal responses while explicitly quantifying whether cluster members share the same underlying trajectory topology.

Clustering operates on the Visit Mean Matrix (VMM), where each molecular feature is represented by its estimated expression across all study visits and experimental groups. Depending on the selected estimator, the VMM may contain empirical visit means, linear mixed-model predictions, generalized additive model predictions, polynomial regression estimates or other visit-level summaries.

For each feature *i*, the longitudinal representation is

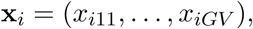

where *G* denotes the number of experimental groups and *V* the number of visits. Features may optionally be standardized before clustering to emphasize trajectory shape rather than absolute expression magnitude.

LongOmicsTraj currently supports centroid-based clustering (k-means), partitioning around medoids (PAM), and model-based clustering using finite mixtures of polynomial regression models through the `lot_from_flexmix_clusters()` function, which interfaces with the `flexmix R` package (version 2.3-20) [11]. For the GLUCOLD analysis, standardized visit mean trajectories from the corticosteroid-withdrawal group were fitted using quadratic regression mixture models. Candidate solutions containing *k* = 2*, . . .,* 5 clusters were evaluated using five random initializations (`nrep` = 5) with a fixed random seed (42). For each candidate solution, LongOmicsTraj computed the Bayesian Information Criterion (BIC), weighted topology purity, the proportion of clusters satisfying a topology-purity threshold (*P_c_ ≥* 0.90), and the minimum cluster size (5 genes). The selected solution was defined as the smallest candidate satisfying both the topology-purity and minimum cluster-size criteria; if no candidate satisfied these constraints, the solution with the lowest BIC was selected. Clustering is performed on continuous longitudinal trajectories, whereas trajectory topology is subsequently evaluated using Ordinal Trajectory States (OTS).

#### 5.3.1 Cluster trajectory topology

Following clustering, each cluster centre is converted into adjacent temporal changes, thresholded, and assigned an OTS exactly as described for individual molecular features. The resulting cluster topology represents the dominant temporal behaviour of the cluster centroid.

In addition, every molecular feature retains its own feature-level OTS assignment. Consequently, each cluster contains a distribution of feature-level topologies rather than a single trajectory.

#### 5.3.2 Topology purity

Cluster homogeneity is quantified using topology purity.

For cluster *c*, let

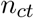

denote the number of molecular features assigned topology *t*, and let

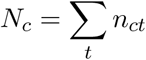

be the cluster size.

The dominant topology is

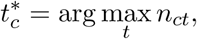

and topology purity is defined as

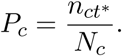

Purity therefore measures the proportion of cluster members sharing the most frequent topology. Values close to one indicate that the cluster represents a single longitudinal behaviour, whereas lower values indicate mixtures of multiple trajectory topologies.

#### 5.3.3 Topology entropy

Purity summarizes only the dominant trajectory. To quantify overall heterogeneity, LongOmicsTraj additionally computes the normalized Shannon entropy

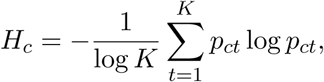

where

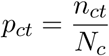

and *K* denotes the number of observed topology classes.

Entropy approaches zero when nearly all members share the same topology and increases as topology diversity increases.

#### 5.3.4 Topology enrichment and depletion

Cluster composition is interpreted relative to the complete clustered dataset rather than in isolation.

For topology *t*,

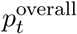

denotes its overall frequency among all clustered molecular features, whereas

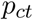

denotes its frequency within cluster *c*. The topology deviation is

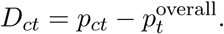

Positive values indicate enrichment of topology *t* within cluster *c*, whereas negative values indicate depletion.

This comparison identifies trajectory patterns that are more or less frequent within a cluster than in the complete clustered dataset.

**Topology-aware model selection** For model-based FlexMix clustering, multiple candidate numbers of clusters are evaluated.

For each candidate *k*, LongOmicsTraj reports: Bayesian Information Criterion (BIC), weighted topology purity, proportion of clusters satisfying a user-defined purity threshold and minimum cluster size.

Rather than relying on likelihood alone, these complementary measures describe both statistical fit and the temporal homogeneity of the resulting clusters. The selected solution is the smallest candidate satisfying the topology-purity and cluster-size criteria. When no candidate satisfies these constraints, the solution with the lowest BIC is selected.

## 6 Results

### 6.1 LongOmicsTraj provides a common topology representation for longitudinal analyses

Figure 1 summarizes the LongOmicsTraj framework. Regardless of the upstream longitudinal estimator used, visit-level trajectories can be converted into a common topology representation through adjacent changes, ordinal state assignment, and Ordinal Trajectory States (OTS). This representation provides a unified language for describing temporal behaviour while preserving the estimated longitudinal trajectory.

Because topology assignment is performed after trajectory estimation, the same downstream queries can be applied consistently across different statistical models, biological representations, and experimental designs. The remainder of this study evaluates the accuracy, robustness, and biological utility of this framework using both simulated and real longitudinal datasets.

### 6.2 Simulation study

#### 6.2.1 Trajectory topology grows exponentially with study duration

The number of possible trajectory topologies increases exponentially with the number of study visits, following 3*^V−^*^1^ (Figure 2A). Consequently, a four-visit study contains 27 distinct trajectory states, increasing to 19,683 states for ten visits. Many of these trajectories share identical start and end points but differ in their intermediate temporal behaviour. Figure 2B illustrates representative Ordinal Trajectory States (OTS), including stable, rebound, transient and oscillatory patterns that cannot be distinguished using endpoint comparisons alone.

**Figure 2:**
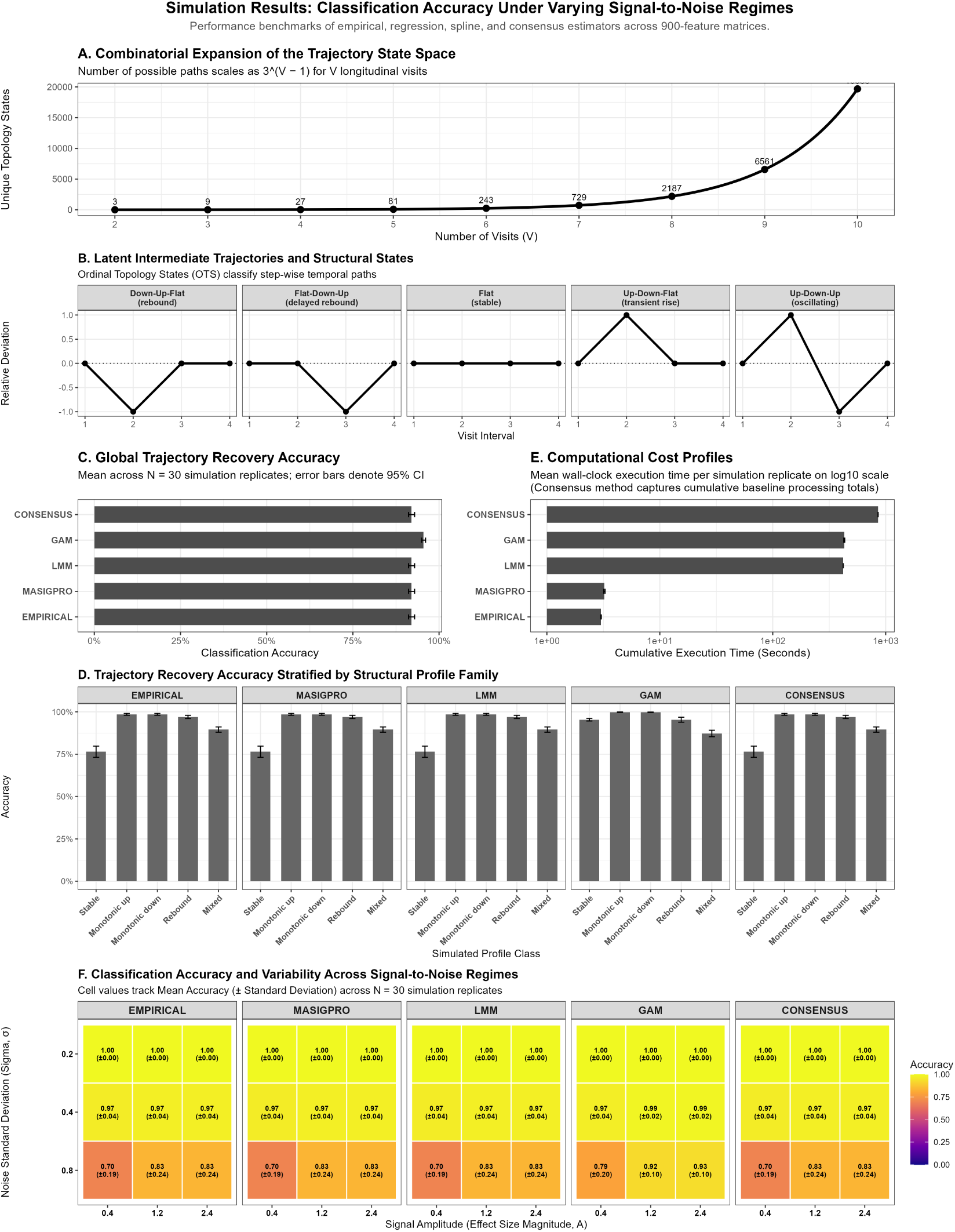
Simulation-based evaluation of LongOmicsTraj across multiple longitudinal estimators. **(A)** Growth of the Ordinal Trajectory State (OTS) space with increasing numbers of visits, showing that the number of possible trajectory topologies scales as 3^(^*^V−^*^1)^. **(B)** Representative OTS describing rebound, delayed rebound, stable, transient, and oscillatory trajectories. **(C)** Overall trajectory recovery accuracy for empirical, maSigPro, linear mixed-effects model (LMM), generalized additive model (GAM), and consensus estimators across 30 simulation replicates (95% confidence intervals). **(D)** Recovery accuracy stratified by simulated trajectory family. **(E)** Mean computational time for each estimator (log_10_ scale), indicating that computational cost is largely determined by the upstream trajectory estimator. **(F)** Classification accuracy across signal amplitudes and noise levels. Cell values report mean accuracy (*±* standard deviation) over 30 simulation replicates. Across all simulations, LongOmicsTraj accurately recovered predefined trajectory topologies, with performance differences primarily 17 reflecting the upstream trajectory estimator rather than the topology representation itself.

#### 6.2.2 Trajectory recovery across longitudinal estimators

All longitudinal estimators accurately recovered the simulated trajectory topology under the baseline simulation setting (Figure 2C). generalized additive model (GAM) achieved the highest overall classification accuracy (95.5%), while the empirical, linear mixed-effects model (LMM), maSigPro and consensus approaches showed similar performance (92.0%). Thus, topology recovery was high across the estimators under the baseline simulation setting, although estimator-specific differences became more apparent under higher-noise conditions.

Performance varied according to trajectory complexity (Figure 2D). All methods classified monotonic trajectories with near-perfect accuracy, whereas stable and mixed trajectories were more challenging. GAM showed improved recovery of stable trajectories, whereas the remaining estimators performed similarly across the other topology classes.

#### 6.2.3 Computational efficiency

Computation time reflected the complexity of the upstream estimator rather than LongOmicsTraj itself (Figure 2E). Empirical estimation and polynomial regression completed within a few seconds per simulation replicate, whereas LMM and GAM required several minutes owing to iterative model fitting. The consensus estimator incurred the largest computational cost because it combines the outputs of multiple individual estimators.

#### 6.2.4 Robustness across signal and noise settings

Classification accuracy depended on both the magnitude of the underlying temporal change and the level of residual noise (Figure 2F). We evaluated three effect sizes (*A* = 0.4, 1.2, 2.4) and three noise levels (*σ* = 0.2, 0.4, 0.8), with 30 independent replicates for each combination, giving 270 simulated datasets.

At low noise (*σ* = 0.2), all estimators achieved 100% topology recovery across all three effect sizes. At intermediate noise (*σ* = 0.4), accuracy remained high. The empirical, LMM, polynomial-regression and consensus approaches achieved 96.8% accuracy at *A* = 0.4 and 97.5% at *A* = 1.2 and *A* = 2.4. GAM achieved 97.3%, 98.9% and 99.0% accuracy across the same effect sizes.

The largest reduction in accuracy occurred under high noise (*σ* = 0.8), particularly when the true temporal changes were small. At *A* = 0.4, accuracy was 69.6% for the empirical, LMM, polynomial-regression and consensus approaches and 78.8% for GAM. Increasing the effect size improved recovery. At *A* = 1.2 and *A* = 2.4, the non-GAM approaches achieved 83.4% accuracy, whereas GAM achieved 92.4% and 93.0%, respectively.

These results show that topology recovery is reliable when visit-to-visit changes are sufficiently large relative to background variation, but deteriorates when weak temporal signals are combined with high noise. Differences between estimators were most apparent under these more difficult conditions, indicating that the quality of the upstream trajectory estimates influences downstream topology assignment.

### 6.3 LongOmicsTraj identifies withdrawal-specific corticosteroid response trajectories

We next evaluated the biological utility of LongOmicsTraj using the longitudinal GLUCOLD bronchial biopsy transcriptomic study. A topology query was designed to identify genes exhibiting (i) an initial response to inhaled corticosteroid (ICS) treatment, (ii) maintenance of that response during continued ICS therapy, and (iii) reversal following corticosteroid withdrawal. To isolate treatment-specific effects, trajectories displaying comparable rebound behaviour in the placebo arm were excluded.

Sequential application of these topology constraints progressively reduced the candidate search space while increasing biological specificity (Figure 3A). Of the 20,358 measured transcripts, 814 genes (4.0%) exhibited an initial corticosteroid response. Excluding genes with similar placebo trajectories reduced this set to 624 genes (3.1%), and requiring sustained treatment response followed by reversal after withdrawal identified a final set of 168 genes (0.8%). Thus, topology-based filtering reduced the search space by more than two orders of magnitude while retaining genes with highly specific longitudinal behaviour.

**Figure 3:**
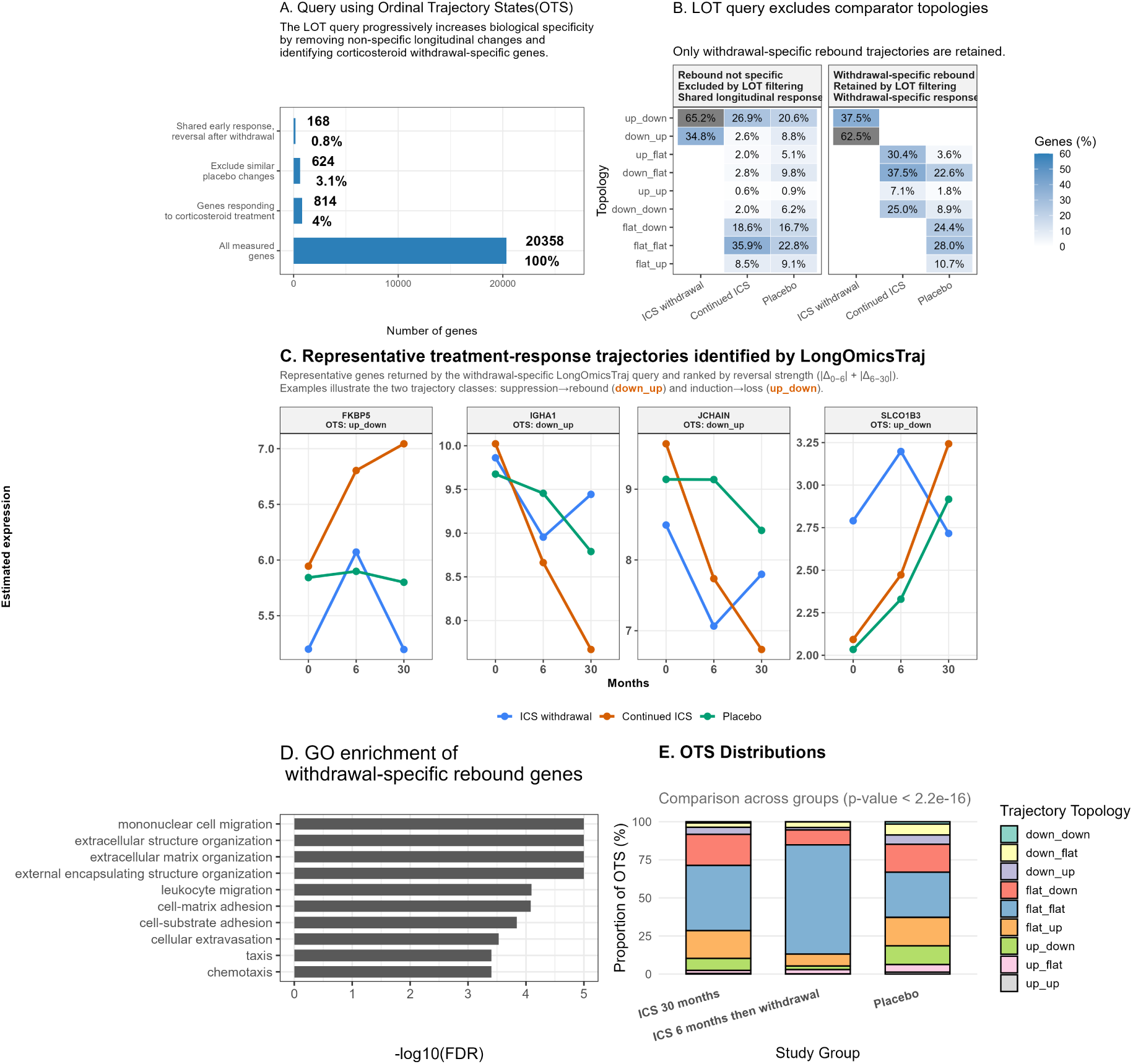
Biological validation of LongOmicsTraj using the GLUCOLD longitudinal corticosteroid intervention study. **(A)** Sequential topology-based queries progressively reduce the candidate gene set by excluding non-specific longitudinal responses and retaining genes exhibiting corticosteroid withdrawal-specific rebound trajectories. **(B)** Distribution of Ordinal Trajectory States (OTS) before and after topology-based filtering, demonstrating selective removal of shared treatment and placebo trajectories while retaining withdrawal-specific rebound responses. **(C)** Representative gene trajectories identified by the withdrawal-specific query, illustrating rebound (down_up) and loss-of-induction (`up_down`) temporal response patterns across treatment groups. **(D)** Gene Ontology enrichment analysis of withdrawal-specific rebound genes, highlighting biological processes associated with immune cell migration, extracellular matrix organization, and cell adhesion. **(E)** Distribution of OTS across treatment groups, demonstrating significant differences in trajectory composition between corticosteroid-treated and placebo groups (*P <* 2.2 × 10*^−^*^16^). Together, these analyses show that LongOmicsTraj identifies biologically coherent, treatment-specific temporal responses while excluding shared or non-specific longitudinal patterns.

Comparator topology filtering was essential for distinguishing treatment-specific responses from non-specific temporal variation (Figure 3B). Before filtering, rebound topologies were observed across all treatment arms, indicating that similar longitudinal patterns can arise independently of corticosteroid withdrawal. Applying the comparator constraints removed these shared responses, leaving only trajectories that rebounded uniquely after withdrawal while remaining absent under continued ICS treatment and placebo. This demonstrates how LongOmicsTraj exploits the complete experimental design rather than analysing treatment arms independently. Representative genes illustrate the temporal behaviours recovered by the query (Figure 3C).

Within each topology class, genes were ranked by a *Reversal Score*, defined as the cumulative magnitude of expression reversal across adjacent transitions,

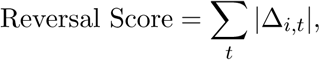

and the two highest-ranking genes were selected for visualization. The `up_down` topology was represented by *FKBP5* and *SLCO1B3*, two well-established steroid response markers that exhibit a characteristic pattern of corticosteroid-driven transactivation followed by a rapid loss of expression upon withdrawal [12, 13]. Conversely, the `down_up` topology was represented by *JCHAIN* and *IGHA1*, which exhibited suppression during treatment followed by recovery after withdrawal. Although these genes differed substantially in baseline expression and response magnitude, they were grouped by their shared temporal topology rather than their absolute expression profiles, illustrating topology-based classification according to temporal pattern rather than absolute expression magnitude.

Functional enrichment analysis of the final withdrawal-specific gene set demonstrated strong biological coherence (Figure 3D). The most significantly enriched Gene Ontology biological processes included extracellular matrix and extracellular structure organization, leukocyte migration, chemotaxis, cell adhesion and cellular migration, consistent with biological processes involved in airway remodelling and inflammation during corticosteroid treatment and withdrawal. Because bronchial biopsies contain a mixture of epithelial, airway smooth muscle, stromal and inflammatory cells, these signals may reflect both cell-intrinsic transcriptional changes and changes in cellular composition. Cell-type deconvolution was not performed in the present analysis. Nevertheless, the functional enrichment supports the biological coherence of the genes identified by topology-based querying rather than serving as an independent validation of their cellular origin.

To examine global differences in longitudinal behaviour, we compared the distribution of OTS across the three treatment arms (Figure 3E). The composition of trajectory topologies differed markedly between continued ICS treatment, corticosteroid withdrawal, and placebo (Pearson’s *χ*^2^ = 8853.5, *df* = 16, *P <* 2.2 × 10*^−^*^16^). Continued ICS treatment and placebo were characterized predominantly by stable trajectories (`flat_flat`), whereas the withdrawal arm showed a relative enrichment of dynamic rebound topologies, including `down_up` and `up_down`, consistent with the expected biological response following corticosteroid withdrawal. These results demonstrate that LongOmicsTraj captures global shifts in longitudinal topology at the transcriptome level in addition to identifying individual treatment-responsive genes.

### 6.4 Topology-aware evaluation of FlexMix trajectory clustering

Figure 4 summarizes the topology-aware evaluation of model-based trajectory clustering in the corticosteroid-withdrawal group.

**Figure 4:**
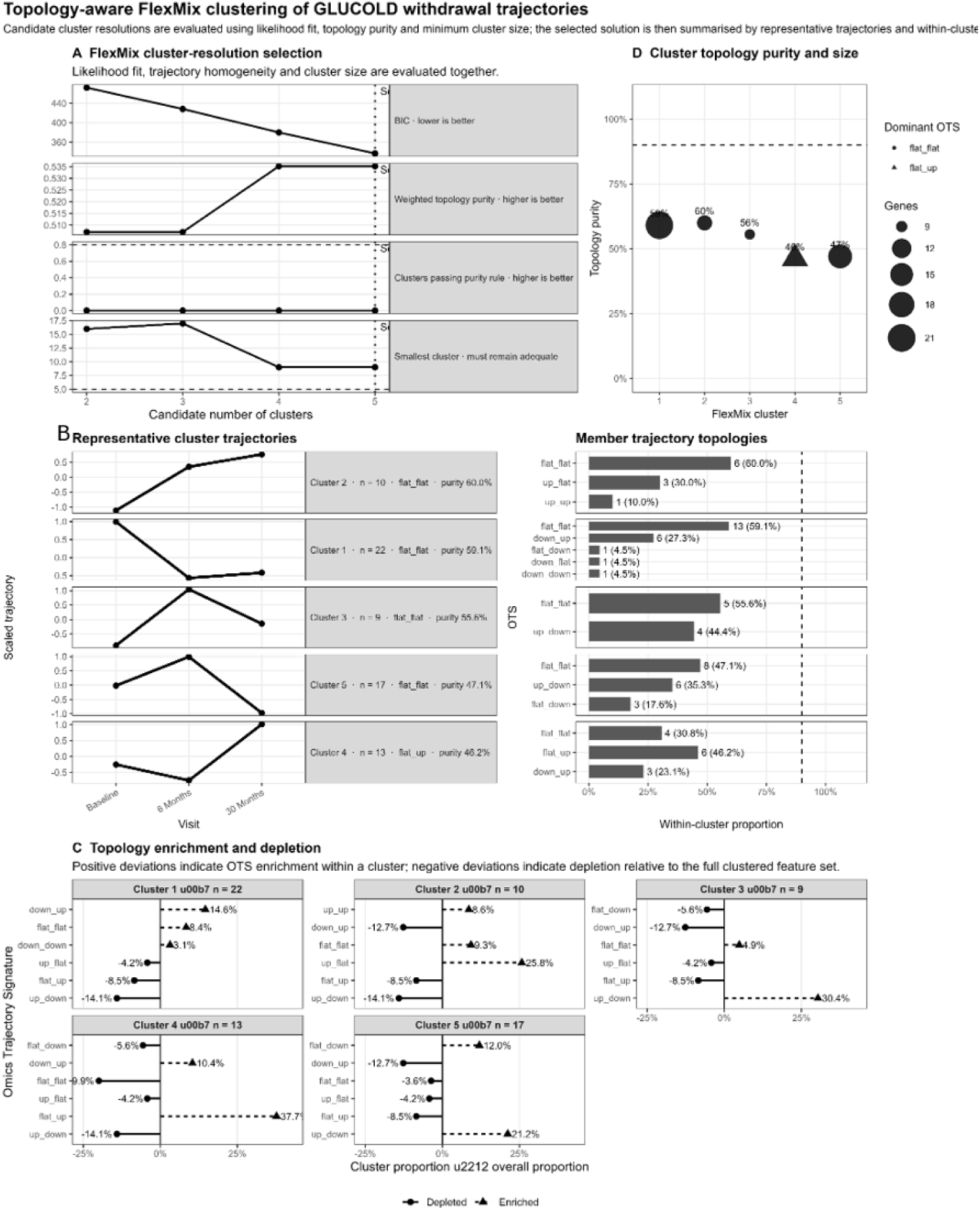
Topology-aware evaluation of FlexMix clustering in the GLUCOLD corticosteroid-withdrawal group. (A) Candidate cluster resolutions evaluated using Bayesian Information Criterion (BIC), weighted topology purity, the proportion of clusters meeting the purity criterion, and minimum cluster size. (B) Representative cluster trajectories and the distribution of feature-level Ordinal Trajectory States (OTS) within each cluster. (C) Enrichment and depletion of OTS relative to the complete clustered dataset. Positive values indicate enrichment and negative values indicate depletion. (D) Cluster size and topology purity. Point size is proportional to cluster size, point shape denotes the dominant OTS, and the dashed line indicates the purity threshold used for model selection.

Candidate models containing between two and five clusters were first evaluated using both statistical and topology-based criteria (Figure 4A). Although BIC decreased monotonically with increasing model complexity, topology-based measures indicated diminishing biological interpretability for larger numbers of clusters. In particular, weighted topology purity remained modest, the proportion of clusters satisfying the purity criterion did not improve, and the minimum cluster size decreased substantially for larger models. These complementary diagnostics show that likelihood-based model selection does not by itself describe the temporal homogeneity of the resulting clusters.

The selected clustering solution identified representative longitudinal response patterns while preserving substantial differences in trajectory topology among cluster members (Figure 4B). The dominant topology of each cluster was readily interpretable from the cluster centre, whereas the accompanying topology distributions quantified the degree of within-cluster heterogeneity. Cluster purities ranged from approximately 46% to 60%, indicating that although dominant temporal behaviours were identifiable, several clusters contained temporally distinct secondary patterns.

Topology-enrichment analysis further characterized these differences (Figure 4C). Rather than simply reporting the most frequent trajectory within each cluster, enrichment analysis compared cluster-specific topology frequencies with their overall prevalence across all clustered genes. Several clusters exhibited clear enrichment for particular rebound or monotonic trajectory classes together with depletion of alternative temporal behaviours, demonstrating that clusters preferentially captured specific dynamic response patterns rather than random mixtures of trajectories.

Finally, Figure 4D summarizes cluster size together with topology purity. Larger clusters were not necessarily more homogeneous, indicating that trajectory homogeneity and cluster size represent complementary characteristics of the clustering solution. Collectively, these results demonstrate that topology-aware diagnostics provide additional information beyond conventional likelihood-based model selection by quantifying how well individual clusters represent coherent longitudinal biological behaviours.

## 7 Discussion

LongOmicsTraj addresses a specific problem in longitudinal omics analysis: how to represent the temporal pattern of a molecular feature after its trajectory has been estimated. It encodes the direction and order of change across visits as an explicit trajectory topology, providing a compact description of temporal behaviour that complements the underlying statistical estimates.

The distinction is particularly relevant for non-monotonic responses. Genes with similar baseline and endpoint expression can follow different intermediate trajectories, including transient activation, delayed response, oscillation or rebound after treatment withdrawal. Existing longitudinal methods, including generalized additive models, maSigPro and ImpulseDE2, can estimate such temporal behaviour [8, 1, 14], but do not generally encode the direction and order of change as a discrete, directly queryable object. LongOmicsTraj adds this representation by mapping visit-level estimates into a finite topology space that can be searched and compared across groups.

An important consequence of separating estimation from representation is estimator flexibility. LongOmicsTraj accepts visit-level estimates from empirical summaries, linear mixed-effects models [7], generalized additive models [8], polynomial regression, or other longitudinal estimators and maps them into the same OTS space. This allows the same topology definitions and downstream queries to be applied across estimation methods without constraining how the trajectories are fitted. In simulations, topology recovery was high across estimators when the longitudinal signal was sufficiently clear, while differences emerged under high-noise conditions. This indicates that the topology framework is compatible with different estimators, while remaining dependent on the quality of the upstream trajectory estimates.

LongOmicsTraj adds a topology-based interpretation layer to conventional longitudinal clustering. Standard clustering groups molecular features according to overall trajectory similarity, but features assigned to the same cluster may still differ in the direction and ordering of their individual temporal changes. LongOmicsTraj retains the continuous trajectories used for clustering and subsequently examines the distribution of feature-level Ordinal Trajectory States (OTS) within each cluster. Topology purity and entropy quantify within-cluster temporal homogeneity, while topology enrichment and depletion describe which trajectory patterns are over-or under-represented relative to the complete clustered dataset. Because topology is evaluated after clustering, these measures can be applied to centroid-, medoid- or model-based clustering without changing the underlying clustering algorithm. They therefore complement conventional criteria such as distance or likelihood by showing whether a statistically defined cluster also represents a coherent temporal pattern.

In the GLUCOLD FlexMix analysis, cluster purities of approximately 46–60% showed that clusters with interpretable average trajectories could nevertheless contain several distinct temporal topologies. This illustrates the added value of topology-aware evaluation: it does not replace clustering or model-selection criteria, but makes within-cluster temporal heterogeneity explicit.

The GLUCOLD analysis illustrates the practical value of making trajectory structure directly queryable. Rather than identifying genes that changed during corticosteroid treatment, the analysis specified a more restrictive temporal pattern: a response that was maintained during continued treatment, reversed after corticosteroid withdrawal, and was not observed under placebo. These comparator-aware constraints substantially reduced the candidate gene set and yielded genes enriched for immune-cell migration, chemotaxis and extracellular-matrix organisation. The resulting trajectories distinguish molecular responses that persist during treatment from those that re-emerge after withdrawal, providing information about treatment durability that is not captured by an endpoint comparison alone. In future longitudinal intervention studies, such patterns could be evaluated as candidate markers of sustained response or relapse and could help formulate hypotheses for treatment-withdrawal or adaptive-treatment designs. These potential clinical applications require prospective validation.

Importantly, the highest-ranked trajectories included established corticosteroid-response genes. *FKBP5* and *SLCO1B3* showed induction during treatment followed by loss after withdrawal, whereas *JCHAIN* and *IGHA1* showed suppression followed by recovery. These patterns are consistent with corticosteroid responses reported independently by Marchi *et al.* [15], supporting the biological relevance of the identified trajectories. The contribution of LongOmicsTraj is not that these genes are inaccessible to conventional longitudinal models, but that their response structure—including induction or suppression, maintenance, and withdrawal-associated reversal—is represented explicitly and can be queried across treatment groups. Features with similar endpoint effects or statistical significance can therefore be distinguished according to the timing and direction of their intermediate changes.

This representation is intended to complement, rather than replace, longitudinal modelling: statistical models provide estimation and inference, while LongOmicsTraj provides a directly queryable description of the temporal pattern represented by those estimates.

### 7.1 Potential clinical applications of topology-based trajectory analysis

A clinically relevant extension of LongOmicsTraj would be to apply the same representation to patient-level trajectories. In longitudinal intervention studies, patients could be characterised according to the timing, persistence and reversal of biological or clinical responses, allowing investigation of heterogeneity in treatment course rather than response magnitude alone. With sufficiently frequent sampling, such analyses could distinguish early from sustained responses and identify trajectories associated with subsequent loss of response or relapse. Whether these temporal patterns provide useful markers of treatment durability or improve patient stratification will require validation against clinical outcomes in independent cohorts.

The representation is also not restricted to transcriptomic measurements. Longitudinal proteomic or metabolomic profiles, cell populations, imaging-derived phenotypes, lung function, FeNO and patient-reported outcomes could, in principle, be represented within the same topology space. This may provide a common framework for comparing the temporal organisation of molecular and clinical measurements within longitudinal studies. Whether such shared representations improve multimodal integration or prediction of clinical outcome remains to be established.

Several limitations should be considered. First, the framework requires a complete Visit Mean Matrix (VMM) but does not itself model missing longitudinal observations. Instead, missing-data handling is delegated to the trajectory-estimation stage. Estimators such as linear mixed-effects models and generalized additive models can accommodate incomplete or unbalanced longitudinal designs under their respective modelling assumptions and can provide complete visit-level predictions for VMM construction. In contrast, empirical trajectory summaries require sufficient observed data at each visit or appropriate preprocessing before topology assignment. This separation preserves flexibility in the choice of upstream estimator without imposing a single missing-data strategy.

Second, OTS assignment depends on the threshold used to distinguish *up*, *down* and *flat* transitions. Features with adjacent changes close to the selected threshold may therefore change topology under alternative parameter settings. LongOmicsTraj provides several thresholding strategies rather than assuming a universally optimal threshold, and sensitivity analysis is recommended when conclusions depend on specific topology assignments.

Third, the current implementation represents trajectory topology through discrete transitions between adjacent visits. This provides an interpretable summary for studies with a small number of clinically meaningful time points, but it necessarily simplifies continuous temporal variation. Future extensions could incorporate uncertainty in visit-level estimates or state assignment, or use continuous trajectory representations for more densely sampled time series. As longitudinal multi-omics studies become increasingly common [16, 17], explicit representation of temporal organisation may complement existing approaches focused on statistical inference, clustering or prediction. We propose that trajectory topology constitutes a complementary molecular phenotype, describing the temporal organisation of molecular responses independently of their absolute magnitude. LongOmicsTraj provides an estimator-agnostic framework for representing, querying and comparing these trajectory topologies across molecular features, biological pathways and longitudinal studies.

